# Phosphorylation on serine 72 modulates Rab7A palmitoylation and retromer recruitment

**DOI:** 10.1101/2024.04.01.587555

**Authors:** Graziana Modica, Laura Tejeda-Valencia, Etienne Sauvageau, Juliette Maes, Olga Skorobogata, Stephane Lefrancois

## Abstract

The small GTPase Rab7A has a key role in regulating membrane trafficking at late endosomes. By interacting with several different effectors, this small GTPase controls late endosome mobility, orchestrates fusion events between late endosomes and lysosomes, and participates in the formation of and regulates the fusion between autophagosomes and lysosomes. Rab7A is also responsible for the spatiotemporal recruitment of retromer, which is required for the endosome-to-TGN retrieval of cargo-receptors such as sortilin and CI-MPR. Recently several post-translational modifications have been shown to modulate Rab7A functions, including palmitoylation, ubiquitination and phosphorylation. Here we show that phosphorylation of Rab7A at serine 72 is important to modulate its interaction with retromer, as the non-phosphorylatable Rab7A_S72A_ mutant is not able to interact with and recruit retromer to late endosomes. We have previously shown that Rab7A palmitoylation is also required for efficient retromer recruitment. We found that palmitoylation of Rab7A_S72A_ is reduced compared to the wild-type protein, suggesting an interplay between S72 phosphorylation and palmitoylation in regulating the Rab7A/retromer interaction. Finally, we identify NEK7 as the kinase required to phosphorylate Rab7A to promote retromer binding and recruitment.

## Introduction

Late endosomes are a highly dynamic compartment where the fate of proteins arriving from different cellular compartments is decided. The small GTPase Rab7A is a principal regulator of trafficking events at late endosomes. Indeed by engaging different effectors, Rab7A can coordinate late endosome-lysosome and autophagosome-lysosome fusion (McEwan et al., 2015), late endosome movement (van der Kant et al., 2013), positioning (Cantalupo et al., 2001; Johansson et al., 2007; Jordens et al., 2001; Rocha et al., 2009), and endosome-to-trans Golgi network (TGN) retrieval of the lysosomal sorting receptors, cation-independent mannose-6- phosphate receptor (CI-MPR) and sortilin (Canuel et al., 2008; Rojas et al., 2008; Seaman et al., 2009).

An increasing amount of data highlights the role of post-translational modifications (PTMs) as a mechanism regulating multiple Rab7A functions (Modica and Lefrancois, 2020). Rab7A is irreversibly prenylated soon after its translation on two C-terminal cysteines (C205, C207), and this PTM is required for its proper membrane anchoring and localization. Indeed Rab7C205,207S mutant fails to bind to endosomal membranes and is almost completely cytosolic (Modica et al., 2017). In addition to prenylation, other modifications such as ubiquitination (Sapmaz et al., 2019; Song et al., 2016), phosphorylation (Francavilla et al., 2016; Heo et al., 2018; Malik et al., 2021; Ritter et al., 2020; Shinde and Maddika, 2016) and palmitoylation (Modica et al., 2017) have been shown to play a role in regulating Rab7A function.

We have previously shown that Rab7A can be palmitoylated on cysteines 83 and 84, and that this reversible modification is required for efficient retromer recruitment and function at endosomes (Modica et al., 2017). Retromer is an evolutionary conserved complex composed of a trimer of vacuolar sorting proteins (Vps) 26, Vps35 and Vps29, that is responsible for the recognition of, and the endosome-to-TGN retrieval of the lysosomal cargo receptors sortilin and CI-MPR (Arighi et al., 2004; Canuel et al., 2008; Seaman, 2004). At the TGN, these receptors recognize and bind soluble lysosomal resident proteins such as cathepsin D and prosaposin, and mediate their trafficking to the endosome via clathrin coated vesicles. Once at the endosome, the more acidic pH of this compartment induces the release of cargo that is eventually trafficked to the lysosome (Bonifacino and Traub, 2003; Coutinho et al., 2012; Luzio et al., 2014). At late endosomes, the receptor is recognized and bound by retromer, and retrieved back to the TGN for another round of trafficking (Arighi et al., 2004; Seaman, 2004). Impaired retromer function results in the accelerated lysosomal degradation of CI-MPR and sortilin and dysfunction of lysosomes (Arighi et al., 2004; Yasa et al., 2020). Rab7A is required for retromer recruitment to endosomes, as down regulation or knock out of this small GTPase results in a significant displacement of retromer from the membrane to the cytosol (Modica et al., 2017; Rojas et al., 2008; Seaman et al., 2009). Palmitoylation regulates the ability of Rab7A to efficiently bind retromer, as in Rab7 knockout (Rab7^KO^) HEK293 cells rescued with non-palmitoylatable Rab7 (Rab7C83,83S), retromer is not efficiently recruited to endosomal membranes (Modica et al., 2017).

Rab7A can be phosphorylated on at least two sites, tyrosine 183 (Y183) and serine 72 (S72). Y183 phosphorylation is mediated by Src kinase, and inhibits the ability of Rab7A to interact with its effector RILP (Lin et al., 2017). LRRK1 has been shown to phosphorylate Rab7 on S72 (Malik et al., 2021), leading to an increased interaction with RILP (Hanafusa et al., 2019), while Tank Binding Kinase 1 (TBK1) also phosphorylates Rab7 on S72, which is necessary for mitophagy (Heo et al., 2018). Transforming growth factor-β (TGF-β)-activated kinase 1 (TAK1) can also phosphorylate Rab7A on this site, and is required for endosomal maturation (Babur et al., 2020).

In this work, we investigate the interplay between phosphorylation of serine 72 and palmitoylation of cysteine 83 and 84 in modulating the ability of Rab7A to interact with retromer. We show that the non-phosphorylatable Rab7A (Rab7A_S72A_) mutant does not efficiently interact with retromer, and fails to rescue retromer membrane recruitment when expressed in Rab7^KO^ HEK293 cells. This phenotype recapitulates the behaviour of non-palmitoylatable Rab7A mutants, and indeed we show that Rab7A_S72A_ is not efficiently palmitoylated. Finally, we identified NIMA-related kinase 7 (NEK7) as a kinase that phosphorylates Rab7A. In HEK293 cells lacking NEK7 (NEK7^KO^), we found significantly less phosphorylated Rab7, decreased Rab7A/retromer interaction, and less membrane bound retromer.

## Results

### Phosphorylation at Serine 72 is required for the Rab7A/retromer interaction

To investigate the role of phosphorylation in modulating the Rab7A/retromer interaction, we first generated Rab7A mutants mimicking a constitutively phosphorylated form (phosphomimetic, Rab7A_S72E_ and Rab7A_Y183E_) or non-phosphorylatable version (phospho-null, Rab7A_S72A_ and Rab7A_Y183F_) of serine 72 or tyrosine 183. Recently, it was shown that Rab7A_S72E_ does not interact properly with the Rab geranylgeranyl transferase (RabGGTase), the enzyme responsible for Rab7A prenylation. This suggests that the protein does not properly localize to endosomal membranes because of the missing prenylated tail (Heo et al., 2018). Therefore, this mutant is likely non-functional, rather than a true phosphomimetic mutant, and was excluded from our study.

To test the effect of phosphorylation on the ability of Rab7A to interact with retromer, we used Bioluminescence Resonance Energy Transfer (BRET). Renilla Luciferase II was fused to the N-terminus of wild-type Rab7A (RlucII-Rab7A) and site directed mutagenesis was used to generate RlucII-Rab7A_S72A_, RlucII-Rab7A_Y183E_ and RlucII-Rab7A_Y183E_. We have previously shown that the addition of the RlucII tag at the N-terminus of Rab7A does not alter the ability of the protein to bind membranes or to rescue retromer recruitment in Rab7A^KO^ HEK293 cells (Modica et al., 2017). We generated BRET titration curves by co-transfecting a constant amount of RlucII-Rab7A, RlucII-Rab7A_S72A_, RlucII-Rab7A_Y183E_ or RlucII-Rab7A_Y183E_ with an increasing amount of Vps26A-GFP10, a subunit of retromer (**Figure 1A**). We have previously shown that Vps26A-GFP10 is integrated into the retromer trimer, and this effector efficiently binds RlucII- Rab7A, but not RlucII-Rab1a, suggesting specificity (Modica et al., 2017; Yasa et al., 2020). By plotting the BRETNET values as a function of the ratio between the fluorescence emission (GFP10 emission) and the luminescence emission (RlucII emission), we calculated the BRET_50_ values from these curves. This value describes the propensity of a protein pair to interact, and the lower the value, the stronger the interaction (Kobayashi et al., 2009; Mercier et al., 2002). RlucII-Rab7A_S72A_ shows a 2.3-fold increase in BRET_50_ compared to wild-type Rab7A (BRET_50_ of 0.0061±0.0011 and 0.0026±0.000512, respectively), suggesting that phosphorylation of S72 is required for the interaction with retromer (**Figure 1B**). While the BRET_50_ values of the RlucII- Rab7A_Y183E_ and RlucII-Rab7A_Y183E_ were slightly higher than wild-type, they were not statistically significant, suggesting this phosphorylation site does not play a role in modulating the Rab7A/ retromer interaction.

**Figure 1.**
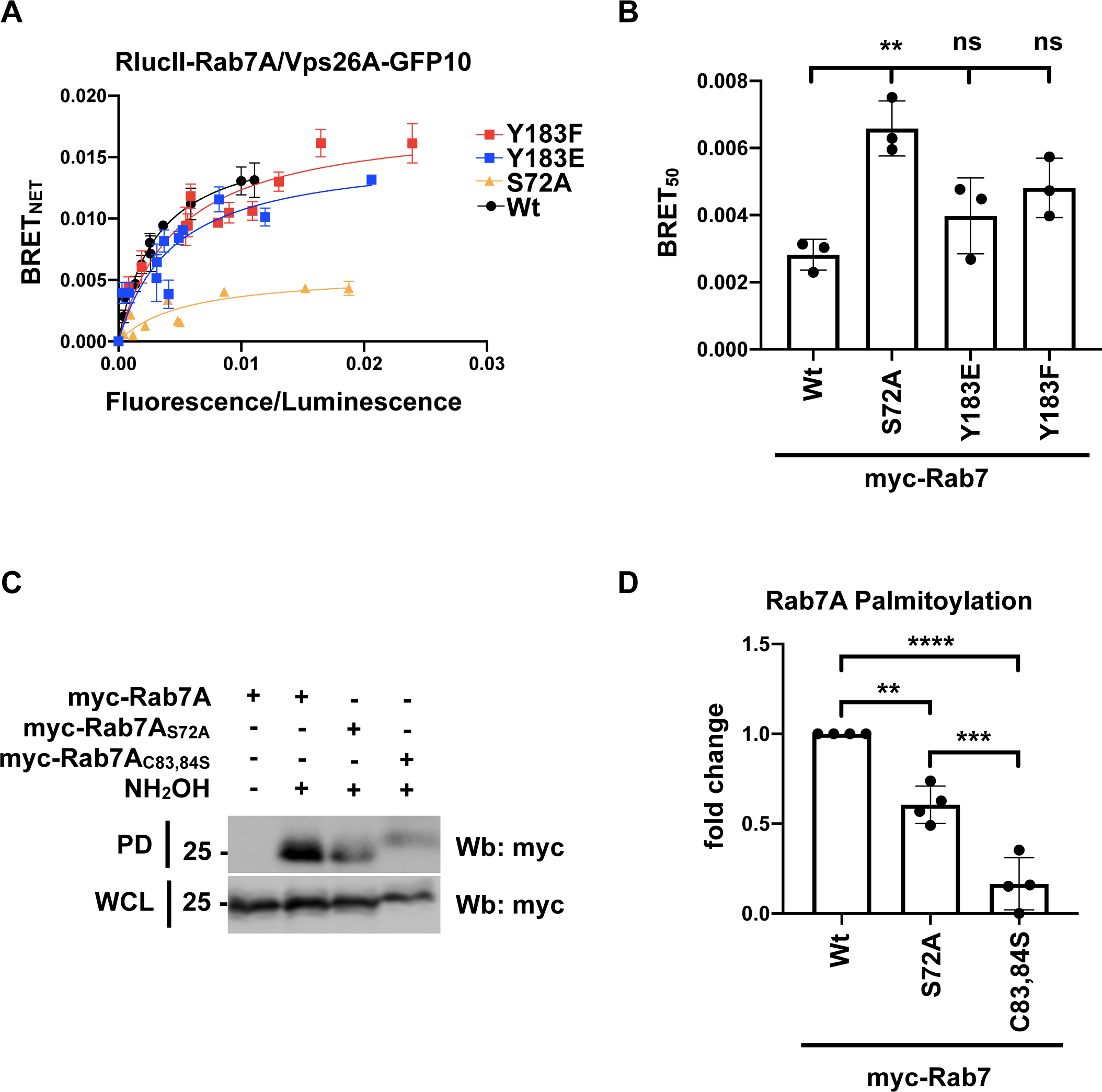
Rab7 S72 phosphorylation regulates Rab7/Retromer interaction. (A) HEK293 cells were transfected with a constant amount of RlucII-Rab7A (black curve), RlucII-Rab7A_S72A_ (yellow curve), RlucII-Rab7A_Y183E_ (blue curve) or RlucII-Rab7A_Y183F_ (red curve), and increasing amounts of Vps26A-GFP10. 48 hours post-transfection BRET analysis was performed. BRET signals are plotted as a function of the ratio between the GFP10 fluorescence over RlucII luminescence. (B) The average of the BRET_50_ extrapolated from the BRET titration curves from 3 separate experiments is shown. Data are represented as mean ± SD. NS, not significant; ** P< 0.01; One-way Anova with Tukey’s post-hoc test. (C) Whole cell lysate (WCL) from HEK293 cells expressing wild-type myc-Rab7, myc-Rab7S72A or myc- Rab7C83,84S were subjected to Acyl-RAC analysis to determine their level of palmitoylation. NH_2_OH: hydroxylamine, PD: pull-down (D) Quantification of 3 separate Acyl-RAC assay experiments. Data are represented as mean ± SD. ** P< 0.01; *** P< 0.001; **** P< 0.0001; One-way Anova with Tukey’s post-hoc test.

We have previously shown that Rab7A palmitoylation on cysteines 83 and 84 is required for optimal retromer recruitment and function at endosomes (Modica et al., 2017). Given that Rab7A_S72A_ is not able to bind to retromer, we wondered if this phenotype could be due to impaired palmitoylation of Rab7A_S72A_. We therefore performed Acyl-Resin Assisted Capture (Acyl-RAC) analysis to compare the levels of palmitoylation between wild-type Rab7A, Rab7A_S72A_ and the non-palmitoylatable mutant, Rab7AC83,84S (**Figure 1C**). We found that Rab7A_S72A_ palmitoylation is significantly reduced compared to wild-type Rab7, but not as significantly as the non-palmitoylatable (Rab7AC83,84S) mutant (**Figure 1D**).

### Rab7A_S72A_ is membrane bound and localized to late endosomes

To determine if the altered Rab7A/retromer interaction we observed was due to changes in the membrane binding and localization of Rab7A, we performed a membrane separation assay as we have previously done (Modica et al., 2017; Yasa et al., 2020). HEK293 cells were transfected with myc-tagged wild-type Rab7A (myc-Rab7A), myc-Rab7A_S72A_ or myc- Rab7A_C205,207S_ (**Figure S1A**). Our membrane separation was successful as the cytosolic protein tubulin was found in the soluble (S) fraction containing the cytosol, while the integral membrane protein Lamp2 was found in the pellet fraction (P) containing the membrane fraction. While myc- Rab7A was membrane bound, as expected, analysis of 3 independent experiments found that the Rab7A prenylation mutant (Rab7A_C205,207S_) was found almost exclusively in the soluble fraction (**Figure S1B**). The non-phosphoratable mutant, Rab7A_S72A_, was found in the pellet fraction (**Figure S1B**). Although Rab7A_S72A_ was membrane bound, we wanted to exclude that the altered interaction of Rab7A_S72A_ with retromer was not due to a defect in the localization of the mutant protein. We co-transfected U2OS cells with the late endosome protein Lamp1- Cerulean and either wild-type myc-Rab7A (**Figure S1C**), myc-Rab7A_S72A_ (**Figure S1D**) or myc- Rab7A_C205,207S_ (**Figure S1E**) and performed co-localization analysis to determine the Pearson’s coefficient. While myc-Rab7A_C205,207S_ had significantly reduced co-localization with Lamp1- Cerulean compared to wild-type myc-Rab7A, myc-Rab7A_S72A_ shows no differences in co- localizing with Lamp1-Cerulean, suggesting that the reduced binding to retromer is not due to an altered localization (**Figure S1F**).

### Phosphorylation on S72 is required for retromer recruitment

Since Rab7A_S72A_ palmitoylation is reduced and unable to interact with retromer efficiently, we next asked if S72 phosphorylation is required to efficiently recruit retromer to endosomes. To answer this question, we performed rescue experiments in our previously generated HEK293 Rab7 knockout (Rab7^KO^) cell line (Modica et al., 2017). We determined the intensity of retromer (Vps26A) using immunofluorescence microscopy and image analysis in wild-type (**Figure 2A**), Rab7A^KO^ cells (**Figure 2B**) or Rab7A^KO^ HEK293 cells expressing either wild-type myc-Rab7A (**Figure 2C, white stars**), myc-Rab7A_S72A_ (**Figure 2D, white stars**), myc- Rab7A_Y183E_ (**Figure 2E, white stars**), myc-Rab7A_Y183F_ (**Figure 2F, white stars)**, or myc- Rab7A_C205,207S_ (**Figure 2G, white stars**). As Rab7A is required for retromer recruitment (Rojas et al., 2008; Seaman et al., 2009), the absence of Rab7A results in the dissociation of retromer from the membrane resulting in a significant decrease of Vps26A puncta in Rab7A^KO^ cells (**Figure 2B and H**) compared to the parental HEK293 cells (**Figure 2A and H**). The expression of wild-type myc-Rab7A (**Figure 2C and H, white stars)**, myc-Rab7A_Y183E_ (**Figure 2E and H, white stars**) and myc-Rab7A_Y183F_ (**Figure 2F and H, white stars**) rescued Vps26A intensity to the same extent as wild-type Rab7A (**Figure 2C and H, white stars**). As expected, the expression of the prenylation mutant, myc-Rab7A_C205,207S_, did not rescue Vps26A intensity (**Figure 2G and H, white stars)**. Expression of myc-Rab7A_S72A_ also did not rescue Vps26A intensity (**Figure 2D and H, white stars**). Combined with the BRET data demonstrating a decreased Rab7A/retromer, these results suggest that Rab7A S72 phosphorylation is required for proper retromer recruitment to endosomes.

**Figure 2.**
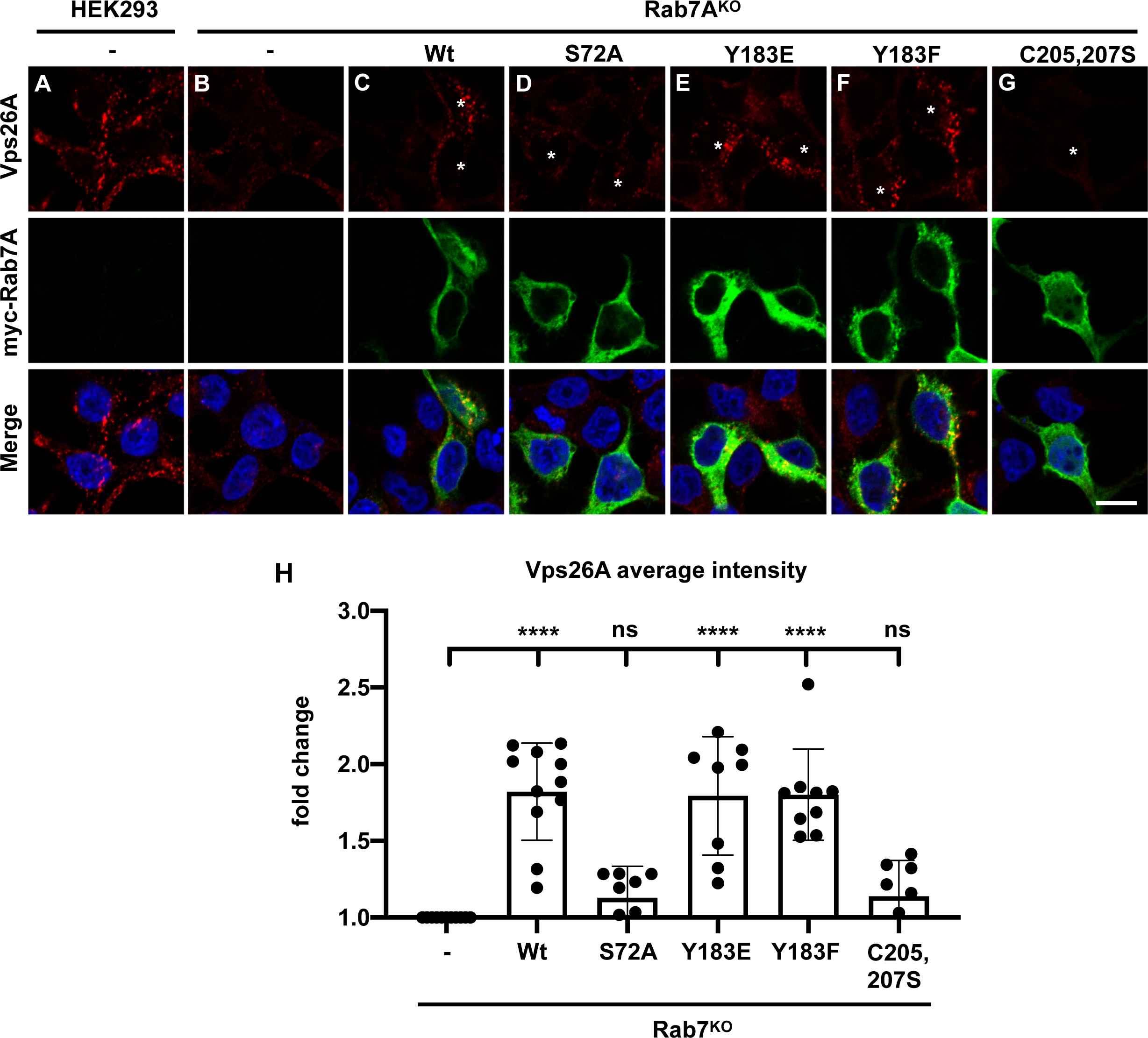
Rab7S72A does not rescue retromer distribution in Rab7^KO^ HEK293 cells. (A - G) Wild-type HEK293 (A), Rab7A^KO^ HEK293 cells (B) or Rab7A^KO^ HEK293 cells expressing wild-type myc-Rab7A (C, stars), myc-Rab7A_S72A_ (D, stars), myc-Rab7A_Y183E_ (E, stars), myc- Rab7A_Y183F_ (F, stars) or myc-Rab7A_C205,207S_ (G, stars) were fixed with 4% PFA and immunostained with anti-Vps26A (red) and anti-myc (green) antibodies. Representative images are shown, scale bar = 10µm. (H) Quantification of Vps26A average intensity from n≥10 cells per condition, values are reported as fold increase compared to Rab7A^KO^. Data are represented as mean ± SD. NS, not significant; **** P < 0.0001; One-way Anova with Tukey’s post-hoc test.

### Inhibiting TBK1 does not alter retromer recruitment

TANK-binding kinase 1 (TBK1) has previously been shown to phosphorylate Rab7A at S72 (Heo et al., 2018). To determine if TBK1-dependent Rab7A phosphorylation plays a role in recruiting retromer, we performed a membrane separation assay in which HEK293 cells were treated with DMSO or with 10 µM of the TBK1 inhibitor MRT67307 for 12 hours (**Figure 3A**). Quantification of 3 independent membrane separation assays found that, compared to DMSO treated cells, retromer (as measured by Vps26A) was slightly more cytosolic in the MRT67307 cells, however, the change in distribution was not statistically significant (**Figure 3B**). Since we observed some changes in retromer distribution in MRT67307 treated cells, we used CRISPR/ Cas9 to generate a TBK1 knockout (TBK1^KO^) HeLa cell line (**Figure S2A**). We then used BRET to test the ability of RlucII-Rab7A and Vps26A-GFP10 to interact in TBK1^KO^ cells (**Figure 3C, red curve**) compared to wild-type cells (**Figure 3C, black curve**). We found no significant changes in the interaction between Rab7A and retromer in TBK1^KO^ cells compared to wild-type HeLa cells (**Figure 3D**). We then used a membrane separation assay to compare the distribution of retromer in TBK1^KO^ and wild-type HeLa cells (**Figure 3E**). We found that the distribution of the retromer subunits Vps26 (**Figure 3F**) is not affected in TBK1^KO^ cells compared to wild-type HeLa cells. Although TBK1 can phosphorylate Rab7A on serine 72, TBK1 phosphorylation is not required for retromer recruitment to membranes.

**Figure 3.**
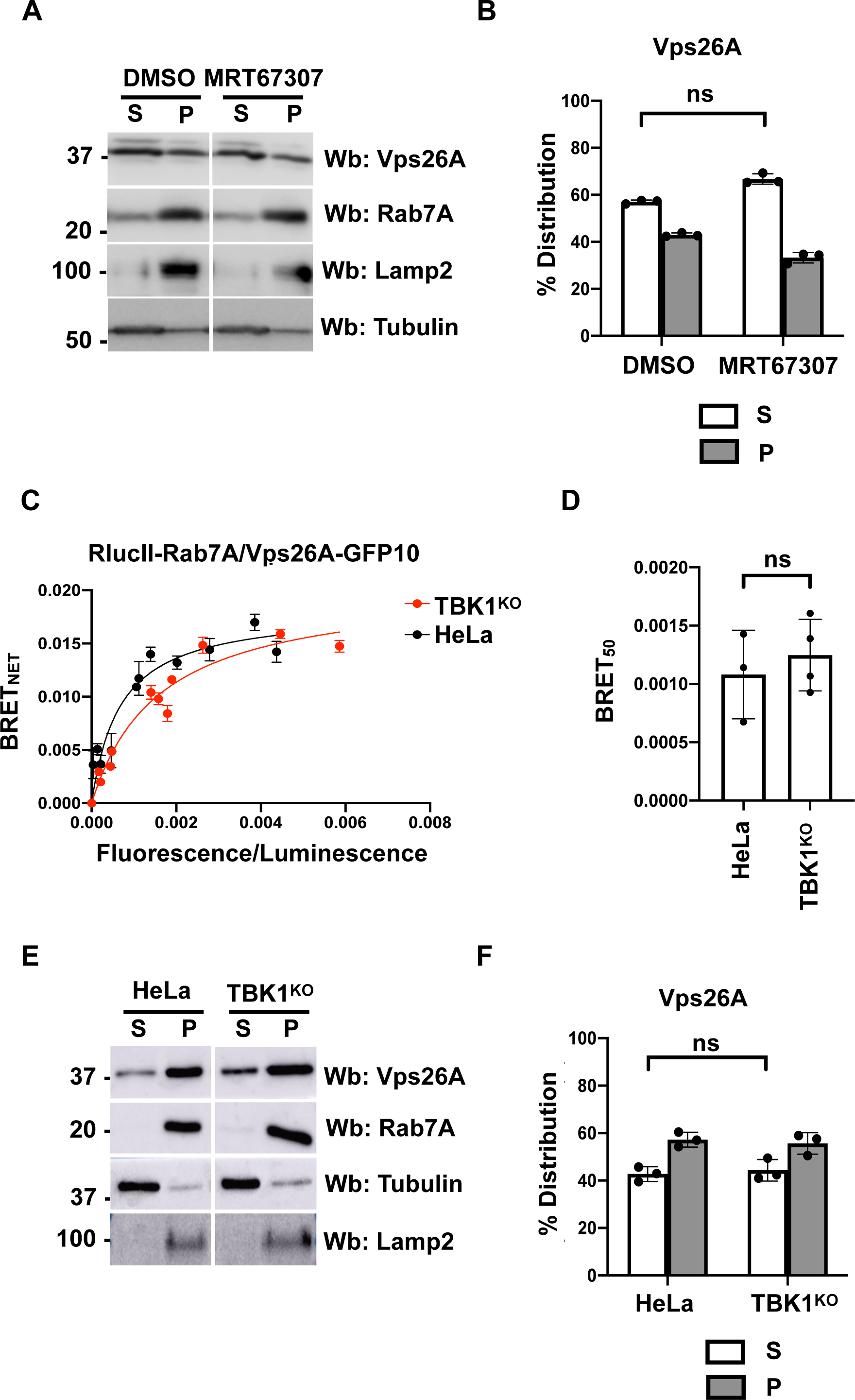
TBK1-dependent Rab7 phosphorylation does not affect retromer recruitment. (A) HEK293 were treated with DMSO or 10µM of the TBK1 inhibitor MRT67307 for 12 hours and subjected to a membrane separation assay. The fractions were subsequently analyzed using Western blot (Wb) with anti-Vps26A, anti-Rab7A, anti-Lamp2 and anti-tubulin antibodies. Lamp2 and tubulin served as markers of the pellet fraction (P) containing membranes, and supernatant fraction (S) containing the cytosol, respectively. (B) Quantification of the distribution of Vps26A from three independent membrane separation assay experiments. Data are represented mean ± SD. NS, not significant; two-tailed unpaired t test. (C) Wild-type (black curve) or TBK1^KO^ (red curve) HeLa cells were transfected with a constant amount of RlucII- Rab7A, and increasing amounts of Vps26A-GFP10. 48 hours post-transfection, BRET analysis was performed. BRET signals are plotted as a function of the ratio between the GFP10 fluorescence over RlucII luminescence. (D) The average of the BRET_50_ extrapolated from the BRET titration curves from 3 separate experiments is shown. Data are represented as mean ± SD. NS, not significant; two-tailed unpaired t test. (E) Wild-type or TBK1^KO^ HeLa cells were subjected to a membrane separation assay. The fractions were subsequently analyzed using Western blot (Wb) with anti-Vps26A, anti-Rab7A, anti-Lamp2 and anti-tubulin antibodies. Lamp2 and tubulin served as markers of the pellet fraction (P) containing membranes, and supernatant fraction (S) containing the cytosol, respectively. (F) Quantification of the distribution of Vps26A from three independent experiments membrane separation assays. Data are represented mean ± SD. NS, not significant; two-tailed unpaired t test.

### Inhibiting TAK1 does not alter retromer recruitment

Recently, TAK1 has been shown to phosphorylate Rab7A (Babur et al., 2020), so we tested if this kinase has an effect in modulating retromer recruitment. We treated HEK293 cells with either DMSO or 10 µM (5Z)-7-oxozeaenol to inhibit TAK1 (**Figure 4A**). Treatment with the TAK1 inhibitor (5Z)-7-oxozeaenol for 7 hours did not affect Rab7A distribution, but had a significant effect on retromer (based on Vps26A), which could be found in the soluble fraction compared to DMSO treated cells (**Figure 4B**). Based on this result, we engineered TAK1 knockout HEK293 cells (TAK1^KO^) using CRISPR/Cas9 (**Figure S2B**). We then used BRET to test the ability of RlucII-Rab7A and Vps26A-GFP10 to interact in TAK1^KO^ cells (**Figure 4C, red curve**) compared to wild-type cells (**Figure 4C, black curve**). We found no significant changes in the interaction between Rab7A and retromer in TBK1^KO^ cells compared to wild-type HeLa cells (**Figure 4D**). We next performed a membrane separation assay to compare the membrane distribution of retromer in wild-type versus TAK1^KO^ HEK293 cells (**Figure 4E**). Quantification of 3 independent experiments found no differences in the membrane distribution of Vps26A (**Figure 4F**) in TAK1^KO^ cells compared to wild-type HEK293 cells.

**Figure 4.**
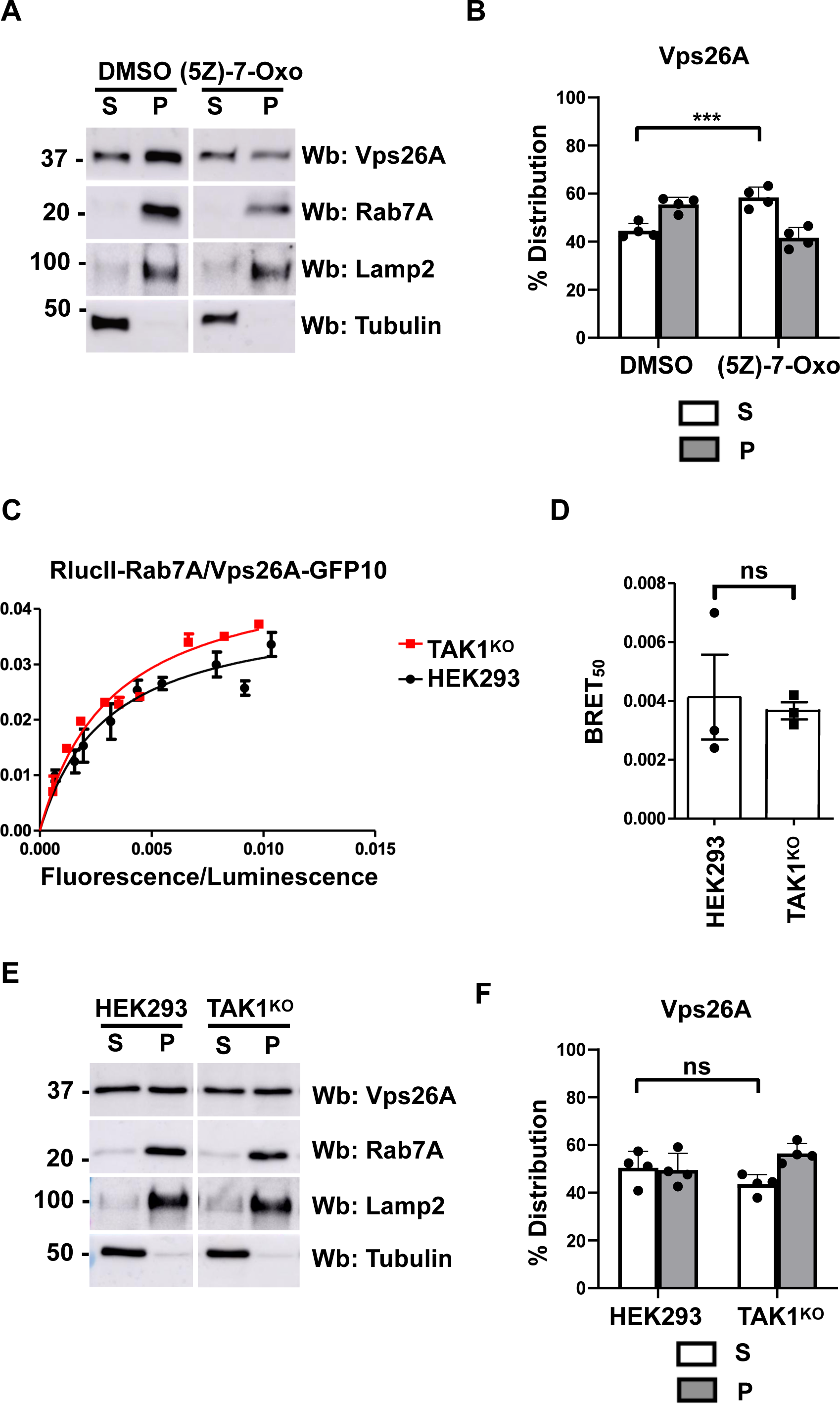
TAK1-dependent Rab7 phosphorylation does not affect retromer recruitment. (A) HEK293 were treated with DMSO or 10µM of the TAK1 inhibitor (5Z)-7-Oxozeaenol for 7 hours and subjected to a membrane separation assay. The fractions were subsequently analyzed by Western blot (Wb) using anti-Vps26A, anti-Rab7A, anti-Lamp2 and anti-tubulin antibodies. Lamp2 and tubulin served as markers of the pellet fraction (P) containing membranes, and supernatant fraction (S) containing the cytosol, respectively. (B) Quantification of the distribution of Vps26A from three independent experiments membrane separation assays. Data are represented as mean ± SD. NS, not significant; *** P< 0.001; two-tailed unpaired t test. (C) Wild-type (black curve) or TAK1^KO^ (red curve) HEK293 cells were transfected with a constant amount of RlucII-Rab7A, and increasing amounts of Vps26A-GFP10. 48 hours post- transfection BRET analysis was performed. BRET signals are plotted as a function of the ratio between the GFP10 fluorescence over RlucII luminescence. (D) The average of the BRET_50_ extrapolated from the BRET titration curves from 3 separate experiments is shown. Data are represented as mean ± SD. NS, not significant; two-tailed unpaired t test. (E) Wild-type or TAK1^KO^ HEK293 cells were subjected to a membrane separation assay. The fractions were subsequently analyzed using Western blot (Wb) with anti-Vps26A, anti-Rab7A, anti-Lamp2 and anti-tubulin antibodies. Lamp2 and tubulin served as markers of the pellet fraction (P) containing membranes, and supernatant fraction (S) containing the cytosol, respectively. (F) Quantification of the distribution of Vps26A from three independent experiments membrane separation assays. Data are represented mean ± SD. NS, not significant; two-tailed unpaired t test.

### NEK7 is required for the Rab7A/retromer interaction

A recent publication demonstrated that knockdown of NEK7 in HeLa cells resulted in the dispersal of CI-MPR into endosomes, suggesting defective retrieval of this sorting receptor (Joseph et al., 2023). This phenotype is similar to the depletion of retromer (Arighi et al., 2004; Seaman, 2004) or Rab7A (Rojas et al., 2008; Seaman et al., 2009). NEK7 is a member of the family of mammalian NIMA-related kinases (NEK proteins) and has been implicated in inflammation (He et al., 2016), and cell cycle regulation (O’Regan and Fry, 2009; Yissachar et al., 2006). As NEK7 has never been associated to phosphorylation of Rab7A, we first tested its role as a Rab7A kinase. As we did not find any commercially available inhibitors of NEK7, we engineered a NEK7 knockout HEK293 cell line (NEK7^KO^) using CRISPR/Cas9 (**Figure S2C**). Using this cell line and a Rab7A phosphorite specific antibody to serine 72 (S72) phosphorylation, we found significantly weaker Rab7A S72 phosphorylation compared to wild-type cells (**Figure 5A**). This decreased phosphorylation was rescued by expressing wild-type HA-NEK7, and partially rescued expressing a putative kinase dead mutant, HA-NEK7_K64M_ (**Figure 5B**). In in vitro assays, NEK7_K64M_ is not able to phosphorylate β-casein (O’Regan and Fry, 2009), however in our hands when expressed in cells, the mutant was able to partially rescue Rab7A S72 phosphorylation, although not as efficiently as wild-type NEK7. We next determined if NEK7 could interact with Rab7A. We attempted co-immunoprecipitation (co-IP) using antibodies to endogenous NEK7 and Rab7A. Although immunoprecipitating with either antibody successfully isolated the target protein, we failed to isolate the other protein. The same negative result was obtained when we attempted the experiment using over expressed and tagged proteins (Data not shown).The transient and potentially weak nature of such interaction might be responsible for our inability to detect via co-IP, But it might be revealed by BRET, as this technique is well suited to detect weak and transient interactions. We generated BRET titration curves by co-transfecting a constant amount of RlucII-Rab7A with an increasing amount of GFP10-NEK7 (**Figure 5C, Black curve**), and we were able to detect an interaction. We also generated BRET titration curves with the non-prenylated mutant, RlucII-Rab7A_C205,207S_ (**Figure 5C, blue curve**), obtaining a linear curve, indicating an absence of interaction. Since this mutant is almost exclusively cytosolic, this data suggests that the Rab7A/NEK7 interaction occurs on the membrane. To confirm the specificity of this interaction we observed by BRET, we performed BRET competition experiments. Cells were transfected with 10 ηg of RlucII-Rab7A and 150 ηg of GFP10-NEK7, and excess amounts of either myc-Rab7A (**Figure 5D, Black curve**) or HA- NEK7 (**Figure 5D, blue curve**). Expressing increasing amounts of either myc-Rab7A and HA- NEK7 inhibited the BRET signal between RlucII-Rab7A and GFP10-NEK7, supporting an interaction between this protein pair. Since NEK7 deletion affected Rab7A phosphorylation and we demonstrated that S72 phosphorylation was required for Rab7A palmitoylation (**Figure 1C and D**), we tested if lack of NEK7 also resulted in decreased Rab7A palmitoylation. Using Acyl-RAC, we determined the level of Rab7A palmitoylation in HEK293 cells, NEK7^KO^ cells, and NEK7^KO^ cells expressing either HA-NEK7 or HA-NEK7_K64M_ (**Figure 5E**). Compared to wild-type HEK293 cells, NEK7^KO^ cells had significantly less palmitoylated Rab7A, which was rescued by expressing HA-NEK7, and partially rescued by expressing HA-NEK7_K64M_ (**Figure 5F**).

**Figure 5.**
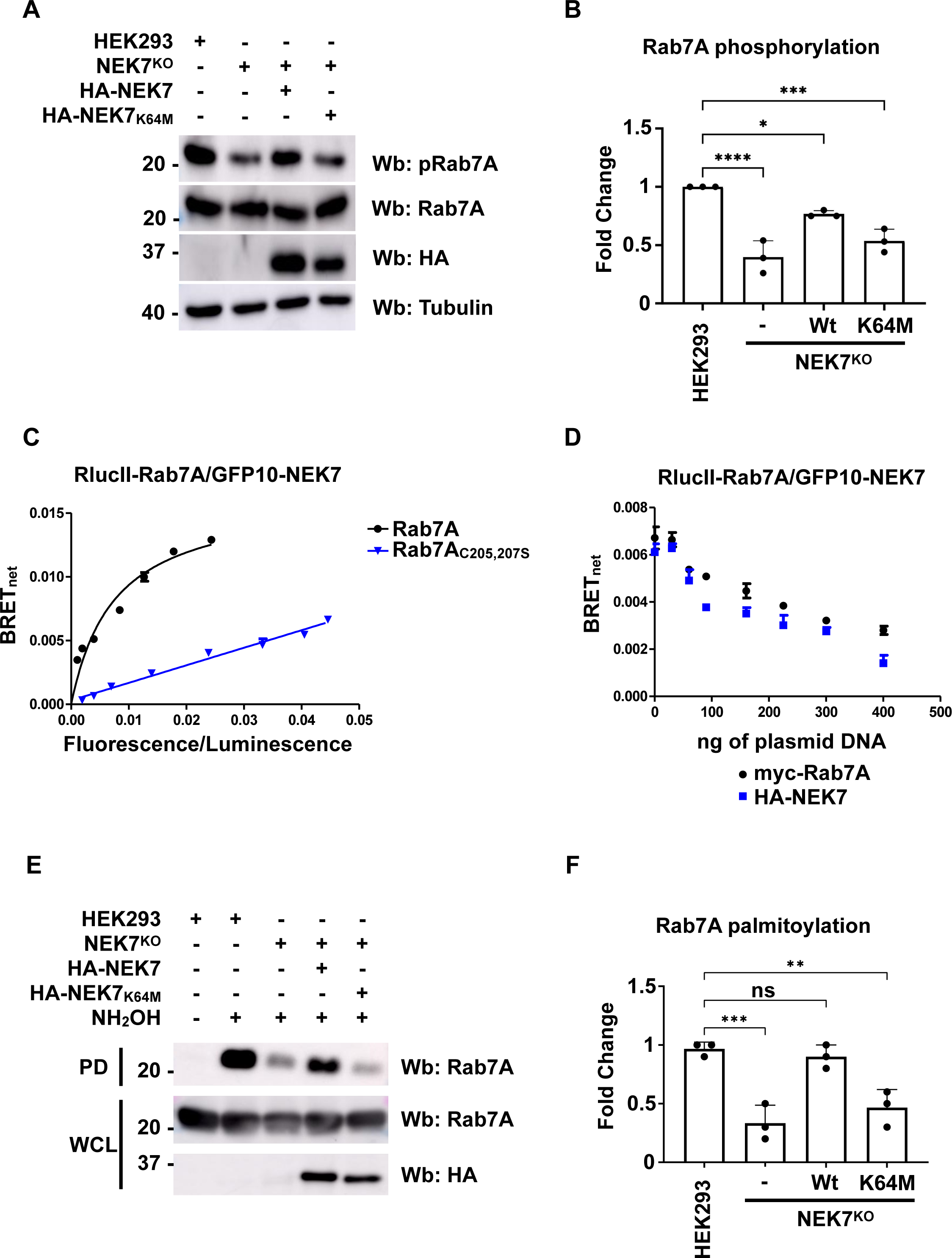
NEK7-dependent Rab7 phosphorylation does not affect retromer recruitment. (A) Lysate from HEK293, NEK7^KO^, and NEK7^KO^ cells expressing either wild-type HA-NEK7 or the kinase dead mutant (HA-NEK7K64M) were analyzed by Western blot (Wb) with anti-Rab7A (phospho S72, pRab7A), anti-Rab7A, anti-HA and anti-actin (as a loading control) antibodies. (B) Quantification of Rab7A serine 72 phosphorylation from 3 independent experiments. Data are represented as mean ± SD. * P< 0.05; *** P< 0.001; **** P< 0.0001; One-way Anova with Tukey’s post-hoc test. (E) Whole cell lysate from HEK293, NEK7^KO^, and NEK7^KO^ cells expressing either wild-type HA-NEK7 or the kinase dead mutant (HA-NEK7K64M) were subjected to Acyl-RAC to determine the level of Rab7A palmitoylation. NH2OH: hydroxylamine, PD: pull- down, WCL: whole cell lysate (F) Quantification of 3 separate Acyl-RAC assay experiments. For the palmitoylation of Rab7A. Data are represented as mean ± SD. NS, not significant, ** P< 0.01; *** P< 0.001; One-way Anova with Tukey’s post-hoc test.

### NEK7 is required for retromer recruitment

We next tested whether NEK7 mediated phosphorylation was required for the Rab7/retromer interaction using BRET (**Figure 6A**). We generated BRET titration curves by expressing a constant amount of RlucII-Rab7A with increasing amounts of Vps26A-GFP10 in wild-type HEK293 cells (**Figure 6A, Black curve**), NEK7^KO^ HEK293 cells (**Figure 6A, blue curve**) or NEK7^KO^ HEK293 cells expressing HA-NEK7 (**Figure 6A, Red curve**). We found 4.36-fold increase in BRET_50_ in NEK7^KO^ HEK293 cells between RlucII-Rab7A and Vps26A-GFP10 in NEK7^KO^ cells compared to wild-type cells (0.0096±0.0039 and 0.0022±0.0009, respectively), suggesting a much weaker interaction (**Figure 6B**), which was rescued by expressing wild-type HA-NEK7 (0.0027±0.0009) (**Figure 6B**). As retromer requires Rab7 for its membrane localization, we tested retromer membrane recruitment in NEK7^KO^ HEK293 cells (**Figure 6C**). We performed a membrane separation assay to compare the distribution of retromer in wild-type HEK293 cells, NEK7^KO^ HEK293 cells and Rab7A^KO^ HEK293 cells (**Figure 6C**). Our membrane separation was successful as the integral membrane protein Lamp1 was found in the pellet fraction (P) that contains membranes, while the cytosolic protein tubules was found in the soluble fraction (S), which contains the cytosol. Quantification from 3 independent experiments showed that compared to wild-type HEK293 cells (44.75% in the pellet fraction), NEK7^KO^ HEK293 cells had significantly less membrane bound retromer (26.25% in the pellet fraction) compared to wild-type cells, but had similar levels to Rab7^KO^ HEK293 cells (28.25% in the pellet fraction (**Figure 6D**). To determine whether retromer recruitment was dependent on NEK7 kinase activity or not, we expressed wild-type HA-NEK7 or the kinase-dead mutant, HA- NEK7K64M, in our NEK7^KO^ HEK293 cells and generated stable cell lines by treating cells with G418. We then performed a membrane separation assay, which was successful as shown by the membrane marker Lamp1 (P), and the cytosolic marker tubulin (S) (**Figure 6E**). Quantification of 3 independent experiments showed that in NEK7^KO^ HEK293 cells expressing HA-NEK7, retromer distribution, as shown by Vps26A Western blotting (Wb), was similar to wild-type HEK293 cells (51.64% in the pellet fraction), while the cells expressing HA-NEK7_K64M_ had a no significant rescue (30.32% in the pellet fraction) (**Figure 6E**).

**Figure 6.**
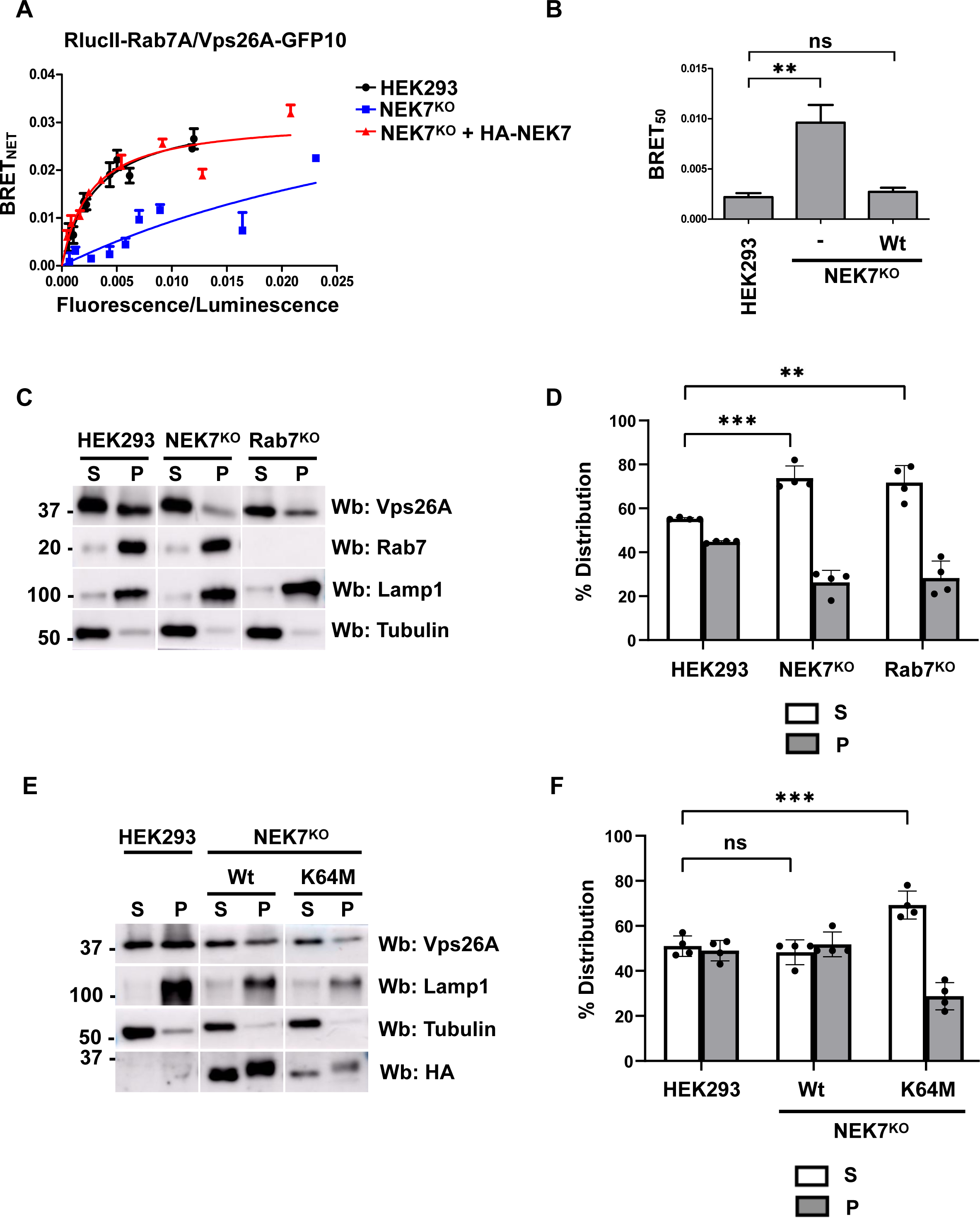
NEK7 is required for the recruitment of retromer. (A) Wild-type (black curve), NEK7^KO^ (blue curve) or NEK7^KO^ expressing HA-NEK7 (red curve) HEK293 cells were transfected with a constant amount of RlucII-Rab7, and increasing amounts of Vps26A-GFP10 to generate BRET titration curves. 48 hours post-transfection BRET analysis was performed. BRET signals are plotted as a function of the ratio between the GFP10 fluorescence over RlucII luminescence. (B) The average of the BRET_50_ extrapolated from the BRET titration curves from 3 separate experiments is shown. Data are represented as mean ± SD. NS, not significant; ** P< 0.01; One-way Anova with Tukey’s post-hoc test. (C) Wild-type, NEK7^KO^ or Rab7^KO^ HEK293 cells were subjected to a membrane separation assay. The fractions were subsequently analyzed using Western blot (Wb) with anti-Vps26A, anti-Rab7, and anti-Lamp1 antibodies. Lamp1 served as a marker of the pellet fraction (P) containing membranes. (D) Quantification of the distribution of Vps26A from three independent membrane separation experiments. Data are represented mean ± SD. NS, not significant; ** P< 0.01; Two- way Anova with Dunnett’s post-hoc test. (E) NEK7^KO^ HEK293 cells stably expressing either HA- NEK7 or HA-NEK7_K64M_ were subjected to a membrane separation assay. The fractions were subsequently analyzed using Western blotting (Wb) with anti-Vps26A, anti-Rab7, anti-Lamp1 and anti-tubulin antibodies. Lamp1 served as a marker of the pellet fraction (P) containing membranes, while tubulin served as the marker for the soluble fraction (S). (F) Quantification of the distribution of Vps26A from three independent membrane separation experiments. Data are represented as mean ± SD. NS, not significant; ** P< 0.01; Two-way Anova with Dunnett’s post- hoc test.

### NEK7 is required to retrieve the lysosomal sorting receptors and lysosomal function

On the endosomal membrane, retromer can interact with the lysosomal sorting receptor sortilin (Canuel et al., 2008). Since we found less membrane bound retromer in NEK7^KO^ HEK293 cells, we tested if the retromer/sortilin interaction would be affected in these cells. We generated BRET titration curves by expressing a constant amount of sortilin tagged to Luciferase (sortilin- RlucII), with increasing amounts of Vps26A-GFP10 in wild-type HEK293 cells (**Figure 7A, black curve**), NEK7^KO^ HEK293 cells (**Figure 7A, blue curve**) or NEK7^KO^ HEK293 cells expressing either HA-NEK7 (**Figure 7A, red curve**). Quantification of 3 independent experiments found a significantly reduced retromer/sortilin interaction in NEK7^KO^ HEK293 cells compared to wild-type HEK293 cells (0.0059±0.0013 versus 0.0021±0.0002, respectively), which was rescued by expressing HA-NEK7 (0.0024±0.0001) (**Figure 7B**). Since retromer is not interacting efficiently with sortilin in NEK7^KO^ HEK293 cells, we would predict decrease retrieval of this cargo protein, and hence more sortilin in endolysosomes compared to wild-type HEK293 cells. To test this, we transfected wild-type, NEK7^KO^ and Rab7A^KO^ HEK293 cells with Luciferase tagged sortilin (sortilin-RlucII) and an endolysosome resident protein, Lamp1, tagged with YFP for energy transfer (YPet), a fluorescent protein derived from Venus (Nguyen and Daugherty, 2005) (**Figure 7C**). Compared to wild-type cells, the BRETnet signal from both NEK7^KO^ and Rab7A^KO^ HEK293 was significantly higher than wild-type cells (**Figure 7D**). This is likely due to more sortilin being retained in endolysosomes, resulting in increased BRET signals. If sortilin is not able to efficiently be retrieved to the TGN for subsequent rounds of sorting, lysosomal activity will be disrupted. We tested the activity of cathepsin L using a fluorogenic substrate. The fluorescence is quenched, until the enzyme, in this case cathepsin L, cleaves the substrate releasing light. As such, a stronger fluorescence signal is interpreted as higher enzymatic activity. Compared to wild-type cells, NEK7^KO^ cells had a 30.34% decrease in cathepsin L activity, which was restored by expressing the HA-NEK7 (95.1% activity compared to wild-type cells), but not HA-NEK7_K64M_ (29.34% decrease). Rab7A^KO^ HEK293 cells were used as a control, and had a similar reduction in cathepsin L activity (22.67% decrease) as NEK7^KO^ cells (**Figure 7D**).

**Figure 7.**
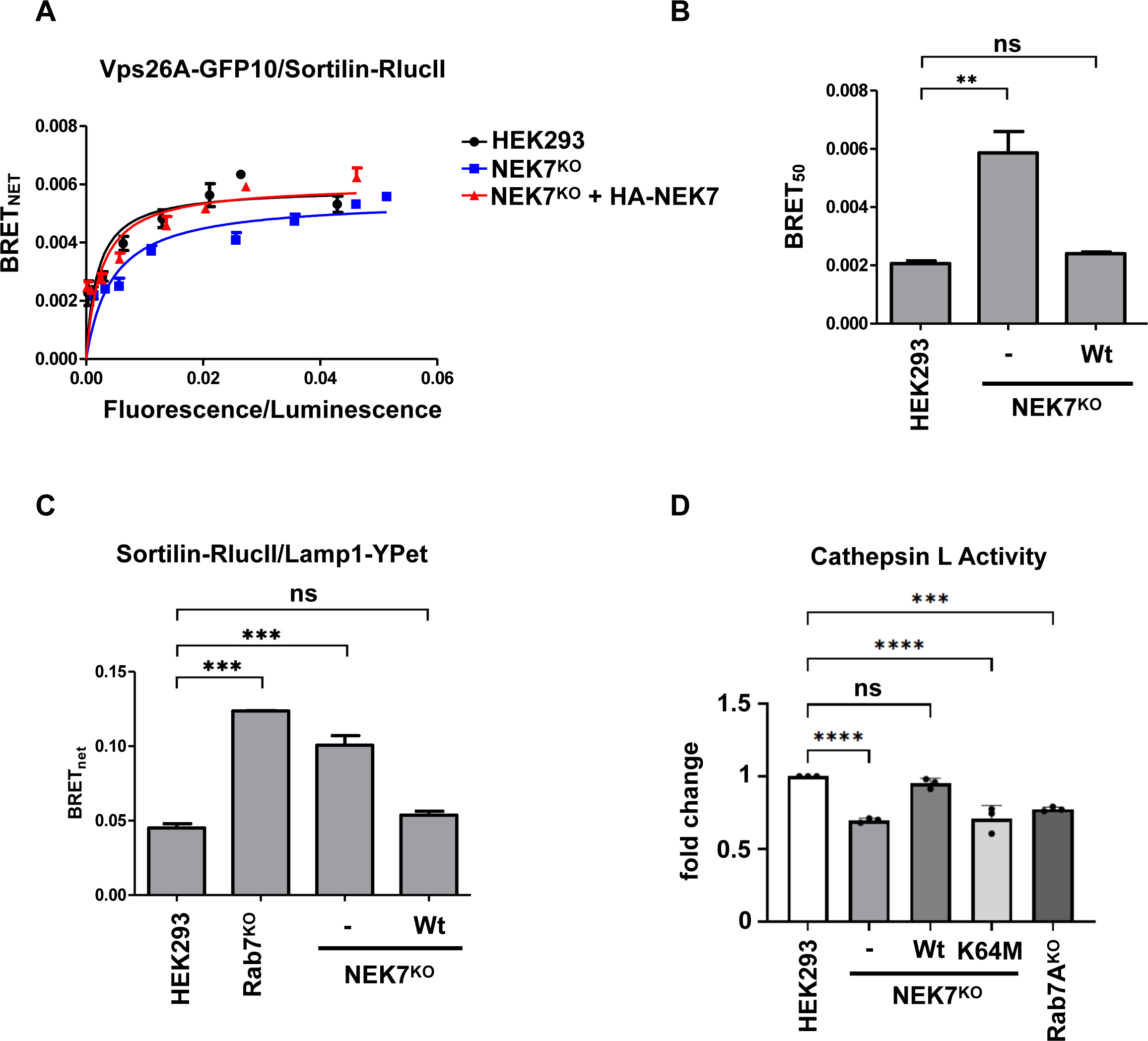
NEK7 is necessary for efficient lysosomal function. (A) Wild-type (black curve), NEK7^KO^ (blue curve) or NEK7^KO^ expressing HA-NEK7 (red curve) HEK293 cells were transfected with a constant amount of sortilin-RlucII, and increasing amounts of Vps26A-GFP10 to generate BRET titration curves. 48 hours post-transfection BRET analysis was performed. BRET signals are plotted as a function of the ratio between the GFP10 fluorescence over RlucII luminescence. (B) The average of the BRET_50_ extrapolated from the BRET titration curves from 3 separate experiments is shown. Data are represented as mean ± SD. NS, not significant; ** P< 0.01; One-way Anova with Tukey’s post-hoc test. (C) HEK293, NEK7^KO^, Rab7A^KO^ cells were transfected with Lamp1-YPet and sortilin-RlucII. Quantification of the BRETnet from 3 independent experiments is shown. Data are represented as mean ± SD. NS, not significant; *** P< 0.001; One-way Anova with Dunnett’s post-hoc. (D) Wild-type, NEK7^KO^, NEK7^KO^ expressing HA-NEK7 or HA-NEK7_K64M_ and Rab7A^KO^ HEK293 cells were incubated with the Cathepsin L MagicRed substrate for 60 mins. Quantification from 3 independent experiments was performed. Data are represented as mean ± SD. NS, not significant; *** P< 0.001; **** P< 0.0001; One-way Anova with Dunnett’s post-hoc test.

## Discussion

Rab7A activity at late endosomes is crucial for several pathways, including late endosome-lysosome and autophagosome-lysosome fusion (McEwan et al., 2015; van der Kant et al., 2013), late endosome movement and positioning (Cantalupo et al., 2001; Johansson et al., 2007; Jordens et al., 2001; Rocha et al., 2009), and late endosome-to-TGN protein retrieval (Rojas et al., 2008; Seaman et al., 2009). The ability of Rab7A to coordinate all these aspects of late endosome physiology is due to the capacity of this small GTPase to interact with different effectors. In this respect, PTMs are able to precisely modulate Rab7A function by favouring interactions with one specific effector according to cellular needs. Our previous work has shown how Rab7A palmitoylation is required to interact with and recruit retromer to endosomes. Here we characterize a further layer of regulation, where the interplay between Rab7A serine 72 phosphorylation and cysteine 83 and 84 palmitoylation is required for Rab7A to efficiently recruit retromer.

### Phosphorylation at Serine 72 does not regulate Rab7 membrane association

Rab7A has been shown to be phosphorylated on at least two sites, tyrosine 183 and serine 72. We and others have shown that phosphorylation on these two sites is not required for Rab7 membrane association. In this respect, the use of the phosphomimetic Rab7A_S72E_ could lead to the wrong interpretation of the role of S72 phosphorylation. Indeed this mutant is mainly localized to the cytosol (Shinde and Maddika, 2016). This result could lead to the conclusion that S72 phosphorylation acts as a switch to terminate Rab7A activity and displace the protein from the membrane. However, previous work showed that Rab7A_S72E_ interacts less with RabGGTase, the enzymes that prenylates Rab7A (Heo et al., 2018). This would suggest that the cytosolic localization of this mutant is due to the absence of the lipid anchor rather than constitutive phosphorylation, resulting in the inability of the small GTPase to stably associate to the membrane.

### NEK7-mediated Rab7 phosphorylation is required for efficient retromer function

TBK1 is related to the family of IKK kinases (I-KB Kinase), and was first identified for its role in promoting the translocation of transcription factors during innate immune response (Abe and Barber, 2014; Bonnard et al., 2000; Cai et al., 2014). In this context, TBK1 also has a role in activating autophagy via the phosphorylation of the autophagic adaptor optineurin (OPTN) for the lysosomal degradation of pathogens (Weidberg and Elazar, 2011). More recently the TBK1- OPTN axis has been described as crucial in coordinating the turnover of damaged mitochondria via mitophagy and hence in maintaining cellular homeostasis (He et al., 2017; Heo et al., 2015; Richter et al., 2016). During mitophagy, TBK1 and Rab7A are independently recruited to the Mitochondrial Outer Membrane (MOM). In this context, TBK1 can phosphorylate Rab7A on serine 72 and this modification is required for the recruitment of ATG9+ membranes for the formation of the autophagosome (Heo et al., 2018). However, our data suggest that, although phosphorylation at serine 72 is required for retromer recruitment, TBK1 is not implicated in this process. TAK1, a meditator of signal transduction in response to TGF-β, has also been shown to phosphorylate Rab7A at serine 72 (Babur et al., 2020). TAK1 has been found to have several roles including playing critical roles in innate immunity. Although it has been shown to phosphorylate Rab7A, it is not implicated in retromer recruitment. Recent work showed that in cells lacking NEK7, CI-MPR was distributed to punctate structures, rather than being primarily localized to the Golgi apparatus (Joseph et al., 2023). This led us to investigate the role of NEK7 in Rab7A serine 72 phosphorylation. We found that this kinase could interact with Rab7A and regulate Rab7A S72 phosphorylation, although we did not demonstrate direct activity. None the less, our data show that this kinase is required for the Rab7A/retromer interaction and retromer recruitment. How this NEK7 is implicated in this process, while TBK1 and TAK1 are not, remains to be elucidates. Could the specific subcellular localization of the kinases themselves play a role?

### Serine 72 phosphorylation is required for efficient Rab7 palmitoylation

The modulation of a protein activity via the combination of several PTMs has been shown previously for several proteins, including member of Ras GTPase superfamily (Liu et al., 2012). Phosphorylation and palmitoylation are two major reversible post-translational modifications used by cells to modulate the activity of proteins according to cellular needs. These PTMs can work in synergy or can have opposite effects in the regulation of a protein (Charych et al., 2010; Gauthier-Kemper et al., 2014; Tian et al., 2008). As for Rab7A, we found a cooperative action of serine phosphorylation and palmitoylation in modulating the ability of this small GTPase to interact with, and recruit retromer. Our data suggest that phosphorylation on S72 is required for efficient palmitoylation, indeed the non-phosphorylatable Rab7A_S72A_ is substantially less palmitoylated compared to wild-type Rab7A. Further supporting the need for phosphorylation for palmitoylation, Rab7A palmitoylation is significantly decreased in NEK7^KO^ HEK293 cells. According to our findings, we can speculate that serine phosphorylation facilitates the interaction of Rab7A with the palmitoylation machinery, possibly by modifying the conformation of the protein itself to favour the interaction with a still unidentified palmitoyltransferase responsible for the addition of the palmitate chain, or by preventing it interaction with thioesterases, which remove the palmitate group. This decreased palmitoylation results in less efficient interaction with retromer, and could explain the inability of Rab7S72A to rescue retromer endosomal recruitment in Rab7^KO^ HEK293 cells.

In summary, we found NEK7-dependent Rab7A s72 phosphorylation as crucial regulator of Rab7/retromer interaction and endosome-to-TGN trafficking pathway. Indeed, in the absence of functional NEK7 activity, the missing S72 phosphorylation hampers efficient Rab7 palmitoylation, leading to decrease retromer recruitment, inefficient retrieval of the lysosomal sorting receptors, and eventually lysosomal dysfunction.

## Materials and Methods

### Reagents, Cloning and mutagenesis

Unless otherwise stated, all reagents used in this study were bought from Fisher Scientific (Ottawa, ON). myc-Rab7A, myc-Rab7AC83,84S, myc-Rab7A_C205,207S_, RlucII-Rab7A, RlucII-Rab7AC83,84S, RlucII-Rab7A_C205,207S_, sortilin-YFP and Vps26A-GFP10 were previously described (Modica et al., 2017; Yasa et al., 2020). myc-Rab7A_S72A_, RlucII-Rab7A_S72A_, myc-Rab7A_Y183E_, myc-Rab7A_Y183F_, RlucII-Rab7A_Y183E_, and RlucII-Rab7A_Y183F_ were generated using site generated mutagenesis. Sortilin-RlucII was generating by cloning the PCR fragment obtained from Sortilin-YFP (a generous gift from Makoto Kanzaki, Tohoku University) into pcDNA3.1Hygro(+)GFP10-RlucII-st2 plasmid (a generous gift from Michel Bouvier, Université de Montreal). pcDNA3-N-HA-NEK7 and pcDNA3-N-HA-NEK7_K64M_ were gifts from Bruce Beutler (Addgene plasmid # 75142 ; http://n2t.net/addgene:75142 ; RRID:Addgene_75142) and (Addgene plasmid # 75143 ; http://n2t.net/addgene:75143 ; RRID:Addgene_75143). mCerulean-Lysosomes-20 was a gift from Michael Davidson (Addgene plasmid # 55382 ; http://n2t.net/addgene:55382 ; RRID:Addgene_55382). YPet-Lysosomes-20 was a gift from Michael Davidson (Addgene plasmid # 56636 ; http://n2t.net/addgene:56636 ; RRID:Addgene_56636). Restriction enzymes used in this study were purchased from New England Biolabs (Danvers, MA). All the mutants described in this work were generated via PCR mutagenesis using cloned PFU polymerase (Agilent Technologies, Santa Clara, CA).

### Antibodies

The following mouse monoclonal antibodies were used: anti-Lamp2 (Wb: 1:500, Abcam ab25631); anti-myc (Wb: 1:1000, IF: 1:500, ThermoFisher Scientific LS132500); anti-HA (Wb: 1 : 1000, Cedarlane Labs 901503); anti-actin (Wb: 1:3000, BD Biosciences 612657); anti-tubulin (Wb: 1:2000, Sigma-Aldrich T9026, St. Louis, MO). The following rabbit monoclonal antibodies were used: anti-Rab7A (Wb: 1:1000, Cell Signalling Technology D95F2); anti-Rab7A (phosphor S72) (Wb: 1:1000, Abcam ab302494), anti-LAMP1 (Wb: 1:1000, Cell Signalling Technology 9091), anti-TAK1 (Wb: 1:1000, Abcam ab109526). The following rabbit polyclonal antibodies were used: anti-Vps26A (Wb: 1:1000, IF: 1:500 Abcam ab23892), anti-TBK1 (Wb: 1:1000, Cell Signalling Technology 3013), anti-NEK7 (Wb: 1:1000, Cell Signalling Technology C34C3).

### Cell culture

All cell lines used in this study were originally obtained from ATCC (Manassas, VA) and regularly screened for contamination. HEK293T, U2OS or HeLa were grown in Dulbecco’s modified Eagle’s medium (DMEM) supplemented with 10% fetal calf serum and penicillin-streptomycin (Thermo Fisher Scientific, Burlington, ON). The Rab7^KO^ HEK293 cell was generated using CRISPR/Cas9 approach as previously described (Modica et al., 2017). The NEK7^KO^ and TAK1^KO^ HEK293 cell lines and the TBK1^KO^ HeLa cell line were generated as previously described (Modica et al., 2017). Transfections were performed with polyethylenimine (PEI) (Fisher Scientific, Ottawa, ON) as previously described (Modica et al., 2017).

### Membrane Separation Assay

Cell pellets were snap frozen in liquid nitrogen and thawed at room temperature (RT). Samples were then resuspended in buffer 1 (0.1 M Mes-NaOH pH 6.5, 1 mM MgAc, 0.5 mM EGTA, 200 µM sodium orthovanadate, 0.2 M sucrose) and centrifuged 5’ at 10000g at 4°C. Supernatant (fraction indicated as S in the text) containing cytosolic proteins was collected and the remaining pellet was resuspended in buffer 2 (50 mM Tris, 150 mM NaCl, 1 mM EDTA, 0.1% SDS, 1% Triton X-100) and centrifuged 5’ at 10000g at 4°C to isolate the supernatant containing membrane proteins (fraction indicated as P in the text). Isolated fractions were analyzed via Western Blot as described in Modica et al., 2017.

### Immunofluorescence

HEK293 and U2OS cell immunofluorescence was performed as described in Dumaresq-Doiron et al., 2010). Immunofluorescence was performed by seeding the cells on coverslip overnight. The following day cells were transfected or treated as indicated in the figure. 24 hrs or 48 hrs after treatment or transfection, coverslips were washed with PBS, fixed with 4% paraformaldehyde (PFA) in PBS for 15 minutes at room temperature (RT). PFA was removed by washing the samples three times with PBS for 5 minutes. Cells were blocked with 0.1% Saponin and 1% BSA in PBS for 1hr at RT followed by incubation with the primary antibody diluted in the blocking solution for 2hrs at RT. Coverslips were washed three times for 5 minutes in PBS and incubated for 1 hour at RT with secondary antibodies conjugated to either AlexaFluor-594 or AlexaFluor-488 in blocking solution. After one wash of 5 minutes in PBS, cells were incubated with DAPI, washed three times for 5 minutes in PBS, mounted on glass slides with Fluoromount G and sealed with nail polish.

### Acyl-RAC

Cells were lysate in TNE (150 mM NaCl, 50 mM Tris, pH 7.5, 2 mM EDTA, 0.5% Triton X-100 and protease inhibitor cocktail) supplemented with 50mM N-Ethylmaleimide (NEM) and incubated for 30 minutes on a rotating wheel at 4°C. Samples were centrifuged 10 minutes at 10000g at 4°C and the collected supernatants were incubated 2 hrs at RT on a rotating wheel. Samples were then precipitated over night with two volumes of cold Acetone at -20°C to remove excess NEM. After washing with cold acetone, the pellet was resuspended in binding buffer (100 mM HEPES, 1 mM EDTA, 1% SDS) with 250 mM hydroxylamine (NH2OH) pH7.5 to cleave palmitate residues off of proteins. Control samples were resuspended in binding buffer containing 250mM NaCl. When the pellet was completely resuspended, Water-swollen thiopropyl sepharose 6B beads (GE Healthcare Life Sciences, Mississauga, ON) were added and samples were incubated 2 hrs at RT on rotating wheel. Beads were then washed 4 times with binding buffer and captured proteins were eluted with 100mM DTT.

### BRET^2^ Assay

HEK293 cells were seeded on 12 well plates overnight followed by transfection with the indicated constructs. 48 hrs after transfections, cells were washed with PBS, detached with 5mM EDTA in PBS and resuspended in 500µl PBS. Samples were then plated in triplicate (90uL/well) on a 96 well plates (VWR Canada, Mississauga, ON). Total fluorescence was measured with an Infinite M1000 Pro plate reader (Tecan Group Ltd., Mannedorf, Switzerland), with the excitation set at 400nm and the emission at 510nm. The Renilla Luciferase substrate coelenterazine 400a (Biotium, Fremont, CA) was added to each well to a final concentration of 5µM and BRET signal was read after 2 minutes incubation at RT. BRET value is calculated as a ratio between the GFP10 emission (500-535nm) over RlucII emission (370-450 nm). To calculate the BRETnet, the BRET obtained by cells expressing only RlucII was subtracted from the BRET value registered from the cells expressing both GFP10 and RlucII. To generate saturation curves, the BRETnet values were plotted as a function of the ratio between the GFP10 signal (Fluorescence) over the RlucII signal (Luminescence).

### Lysosomal Activity

To determine the activity of cathepsin L, wild-type, NEK7^KO^, NEK7^KO^ expressing HA-NEK7 or HA-NEK7_K64M_ and RAB7A^KO^ HEK293 cells were collected and 3 × 10^6^ cells/ml of each cell type were transferred to 96-well black wall plates in triplicate. Cells were then incubated with the Magic Red substrate for 60 min at 37°C protected from light. As cells settled to the bottom, they were gently resuspended by pipetting every 10–20 min to ensure that the Magic Red was evenly dispersed among all cells. The fluorescence intensity of the substrate was measured with a Tecan Infinite M1000 Pro plate reader (Tecan Group Ltd., Mannedorf, Switzerland) with the excitation and emission set at 592 nm and 628 nm, respectively. The average of non-stained sample fluorescence intensities was calculated for each sample and subtracted from the fluorescence reads of the Magic Red-stained samples to eliminate background fluorescence, and signals were standardized using Hoechst stain in each sample.

### Image analysis and statistics

Image analysis was performed using Fiji (Schindelin et al., 2012) and the coloc2 plugins for the co-localization analysis. Statistical analysis was performed using GraphPad Prism Version 8.2.1 (GraphPad Software, San Diego, California USA, www.graphpad.com) and described in the corresponding figure legend.

## Author contribution

Conceptualization, S.L., G.M. Design (Methodology), S.L., G.M., Investigation G.M., O.S., and L.T. Writing - Original Draft, S.L., G.M: Writing - Review & Editing S.L., G.M., L.T., E.S. Funding Acquisition, S.L.

## Acknowledgements

We would like to thank Michel Bouvier (IRIC and Université de Montreal), Regis Grailhe, (Pasteur Institute Korea) for sharing reagents.

---

**Figure S1.**
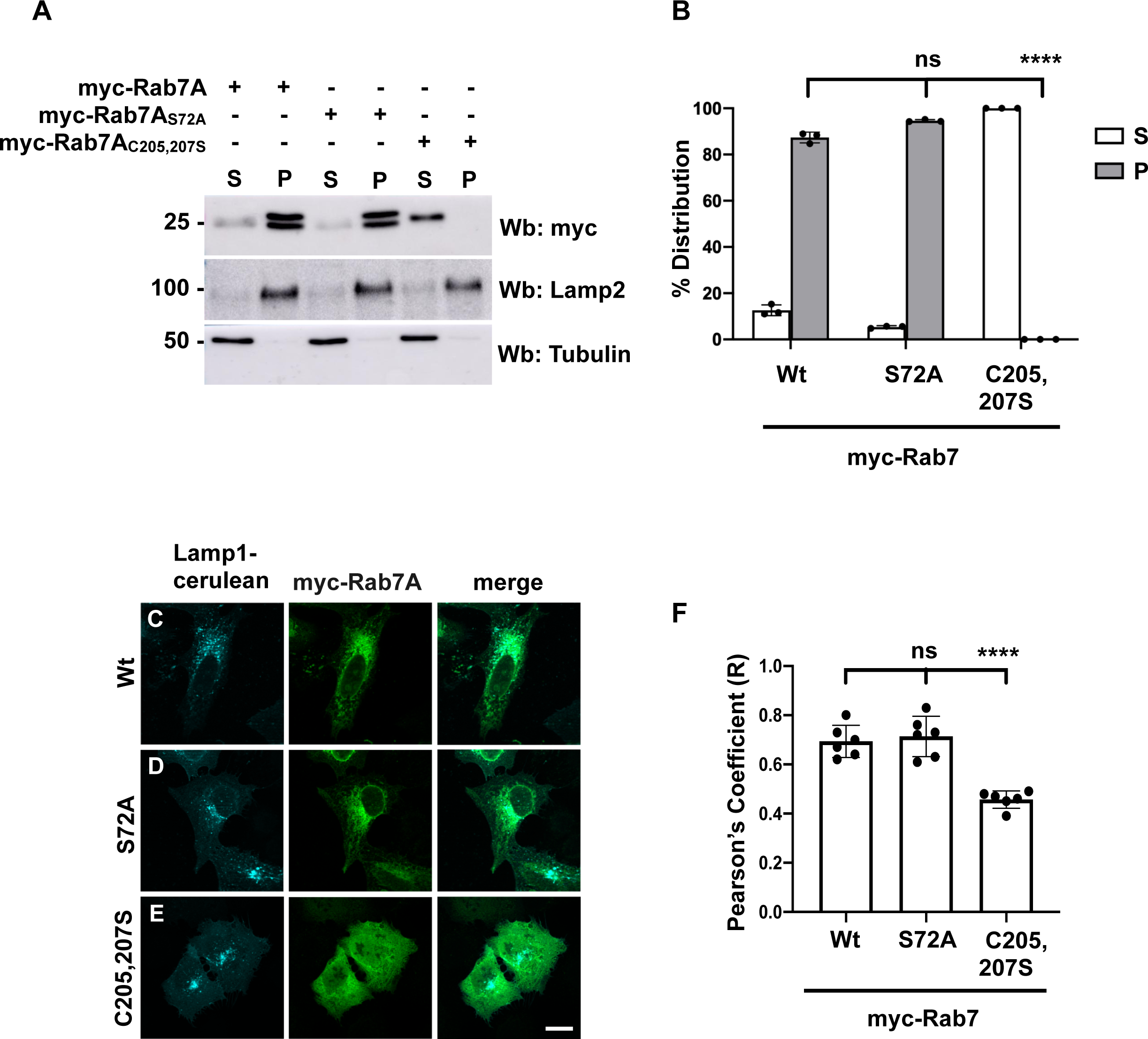
Rab7A S72 phosphorylation is not required for localization. (A) A membrane separation assay was performed on HEK293 cells expressing myc-Rab7, myc- Rab7S72A or myc-Rab7C205,207S. Western blot (Wb) was performed with anti-myc, anti-Lamp2 and anti-tubulin antibodies. Lamp2 and tubulin served as markers of the pellet fraction (P) containing membranes, and supernatant fraction (S) containing the cytosol. (B) Quantification of the distribution of myc-Rab7, myc-Rab7S72A or myc-Rab7C205,207S from 3 independent experiments. Data are represented as mean ± SD. NS, not significant; **** P< 0.0001; One-way Anova with Tukey’s post-hoc test. (C) U2OS cells were co-transfected with Lamp1-cerulean and wild-type myc-Rab7 (C), myc-Rab7S72A (D) or myc-Rab7C83,84S (E). 24 hrs after transfection, cells were fixed with PFA 4% and immunostained with anti-myc antibody (green). Representative images are shown, scale bar = 10µm. (F) Pearson’s correlation coefficient from n≥12 cells per condition. Data are represented as mean ± SD. , n.s. not significant, **** P < 0.0001; One-way Anova with Tukey’s post-hoc test.

**Figure S2.**
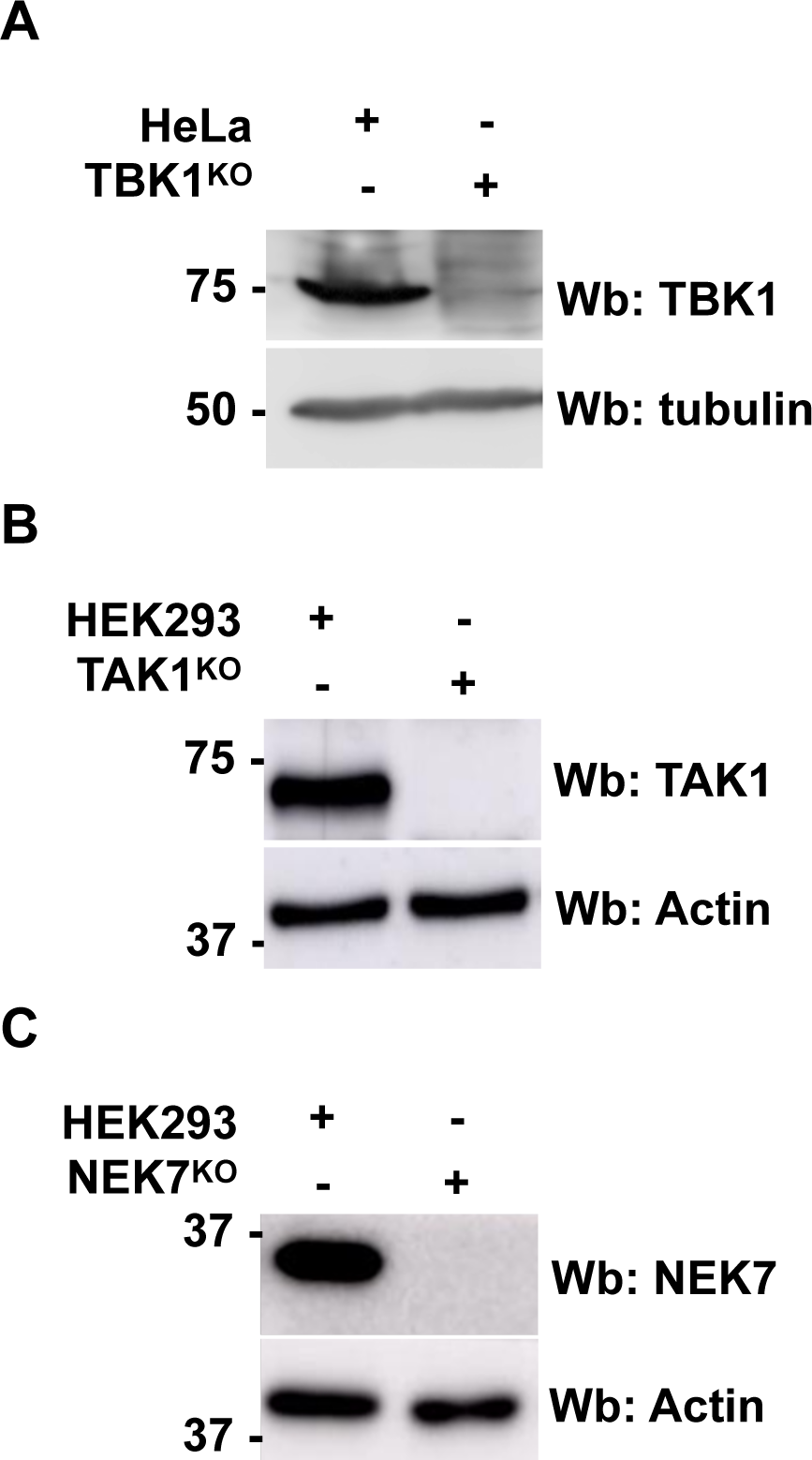
Confirmation of CRISPR/Cas9 knockout. (A) Whole cells lysates from wild-type and TBK1 knockout (TBK1^KO^) HeLa cells was run on a 12% SDS-PAGE gel, transferred to a nitrocellulose membrane, and Western blotting (Wb) was performed using anti-TBK1 and anti-tubulin antibodies. (B) Whole cells lysates from wild-type and TAK1 knockout (TAK1^KO^) HEK293 cells was run on a 12% SDS-PAGE gel, transferred to a nitrocellulose membrane, and Western blotting (Wb) was performed using anti-TAK1 and anti-actin antibodies. (C) Whole cells lysates from wild-type and NEK7 knockout (NEK7^KO^) HEK293 cells was run on a 12% SDS-PAGE gel, transferred to a nitrocellulose membrane, and Western blotting (Wb) was performed using anti-NEK7 and anti-actin antibodies.

